# Alpha-2-Macroglobulin/LRP1 signaling promotes mitochondrial clearance and autophagic extracellular vesicle release in erythroid cells

**DOI:** 10.64898/2026.06.12.731984

**Authors:** J Jacob, SR Castro Pérez, BN Salassa, C Deleschaux, AF Campos Londero, A Dussouchaud, SD Lefevre, GA Chiabrando, MA Ostuni, CM Fader

## Abstract

Despite advances in the understanding of the cellular and molecular mechanisms involved in erythropoiesis, there are still unanswered questions regarding the coordination between autophagy, vesicular trafficking, and endocytic signaling during this process. The complexity of these events suggests the existence of regulatory mechanisms capable of integrating these pathways. In this context, low-density lipoprotein receptor-related protein 1 (LRP1) emerges as a potential modulator given its function as a multifunctional endocytic receptor and its involvement in the regulation of degradation and signaling processes in various cellular models. However, its role in modulating mitophagy, a particular type of autophagy, and its link to vesicular trafficking associated with multivesicular bodies (MVBs) and the release of exosomes during erythroid maturation has been poorly explored. In this regard, alpha-_2_-Macroglobulin (α_2_M), the main physiological ligand of LRP1, has been identified in extracellular vesicles (EVs) in various pathophysiological contexts, suggesting that it may be involved in vesicular dynamics and cellular clearance. In this study, we demonstrate that activated α_2_M (α_2_M*), induces autophagy and particularly mitophagy, in K562 cells, and that LRP1 is directly responsible for this activation. Furthermore, we observed that α_2_M* stimulates the interaction of autophagosomes with MVBs/amphisomes and that EVs from K562 cells are positive for LC3, supporting a close relationship between the endocytic pathway and the autophagic pathway mediated by the α_2_M-LRP1 interaction. Taken together, these findings expand our understanding of erythroid biology and provide a conceptual foundation for exploring altered mechanisms in erythropoietic diseases and for the development of diagnostic and therapeutic strategies.

## Introduction

Hematopoietic progenitors undergo a series of modifications during erythropoietic maturation before achieving their final functional state^1^. These changes include a reduction in cell size, chromatin condensation, production of specific proteins such as hemoglobin, expulsion of the nucleus, loss of mitochondria, ribosomes, and all other organelles through autophagy, and a transition to anaerobic glycolysis^2,3^. Finally, the reticulocyte migrates into the bloodstream, where it completes its differentiation process and transforms into a mature red blood cell^4^. Autophagy is an important process during erythropoiesis allowing cells to remove unnecessary organelles, including mitochondria, ribosomes and the endoplasmic reticulum^1^. This process is characterized by the formation of a double-membraned vesicle called an autophagosome, which engulfs cellular material targeted for degradation^5^. The autophagosome then fuses with the lysosome to produce an autolysosome, which subsequently degrades the vesicular contents^6^. During erythropoiesis, certain proteins that are not required during the maturation stage become sequestered into multivesicular bodies (MVBs) vesicles^4^. With a diameter of around 250–1000 nm, enclosed by a single membrane, MVBs are organelles that contain smaller intraluminal vesicles (ILVs) with a diameter of 50–150 nm^7^. When an MVB fuses with the plasma membrane, these ILVs are released as exosomes into the extracellular environment^8,9^. Exosomes regulate many cellular functions, including differentiation, proliferation, migration, invasion, and apoptosis, in addition to serving as carriers for proteins, microRNAs, and long non-coding RNAs across cells^10,11^. Along with microvesicles and apoptotic bodies, exosomes are part of the wider group known as extracellular vesicles (EVs)^12^. In addition, MVBs can fuse with autophagosomes to form a hybrid vesicle called an amphisome, which also degrades their contents following lysosomal fusion^9^.

Increasing evidence has been reported that abnormalities in erythropoiesis lead to a range of hematological disorders, such as anemias, thalassemia, myelodysplastic syndromes, and leukemias^13,14^. Many of these conditions are associated with abnormalities in autophagy and mitophagy^4^. One example is the deletion of the autophagic genes Bnip3L, Ulk1, Atg7, and FIP200, the loss of which causes mitochondrial retention, defective erythroid differentiation, and anemia^15–17^. In diseases such as β-thalassemia and myelodysplastic syndrome (MDS), abnormal activation of autophagy has been described, contributing to ineffective erythropoiesis and leading to chronic anemia and other hematological disorders^18–20^. Ineffective hematopoiesis, anemia, dissociation between proliferation and differentiation of progenitor cells, and inefficient clearance of aggregated protein are the hallmarks of these types of disorders^21–23^. Other hematological pathologies have also been linked to alterations in autophagic mechanisms, for example, in sickle cell disease (SCD) and systemic lupus erythematosus (SLE), pathogenesis-associated defects in mitophagy have been described^24–26^. Furthermore, accumulation of protein in mature erythrocytes has been documented in SCD, indicating an irregularity in the exosomal process ^27,28^. Additionally, autophagy modulators have been proven to be favorable for both these individuals and those with β-thalassemia^18,24,29^. About other anemias, such as anemia in Pearson’s syndrome, incomplete mitochondrial clearance in reticulocytes has been documented^30^. In leukemias, autophagy plays a dual role, acting both as a survival mechanism for the leukemic clone and as a modulator of therapeutic response^31^. The induction of mitophagy in erythroleukemic cells promotes erythroid differentiation, which could be exploited to reduce malignancy in chronic myeloid leukemias^32^. In acute myeloid leukemia, autophagic flux inhibition enhances sensitivity to chemotherapeutic agents, reducing tumor growth *in vitro* and *in vivo*^33^. Finally, defective autophagy has been connected to chorea-acanthocytosis, a rare neurohematological condition that results in delayed mitochondrial and lysosome degradation^34^. Taken together, these findings demonstrate that autophagy, and more specifically mitophagy, plays a crucial role in erythroid homeostasis and that its alteration is linked to multiple hematological pathologies of different etiology and severity.

Low-density lipoprotein receptor-related protein 1 (LRP1), also known as alpha-_2_-Macroglobulin receptor or differentiation group 91, is a transmembrane protein receptor with numerous cellular and physiological functions that belongs to the low-density lipoprotein receptor superfamily^35^. This receptor is widely expressed on cell surfaces and mediates the endocytosis of over 40 structural and functionally distinct ligands, including proteases, complex inhibitors of proteases, extracellular matrix proteins, growth factors, toxins, and viral proteins^36–38^. Because of its ability to bind multiple ligands, LRP1 plays an important role in the regulation of cell growth and differentiation, cell survival, angiogenesis modulation, cytoskeleton organization, cell migration, and protease-mediated tissue remodeling and inflammation^39^. LRP1 facilitates the ligand internalization through the cytoplasmic tail by clathrin-dependent endocytosis when its extracellular subunit binds to a ligand^35^. LRP1 recycling to the membrane requires ligand-receptor complex dissociation in endosomes^37,40^.

Alpha-_2_-Macroglobulin (α_2_M) is a large plasma glycoprotein that acts as a broad-spectrum protease inhibitor as it can inhibit all families of proteinases, including serine, cysteine, aspartic, and metalloproteinases^41^. In addition, it can actively bind and transport cytokines, hormones, apolipoproteins, growth factors, and misfolded proteins to suppress enzymatic cascades and the kallikrein-kinin system, regulate blood coagulation and fibrinolysis processes, and initiate inflammatory reactions^40^. α_2_M is a 720 kDa glycoprotein formed by assembling four identical 180 kDa subunits^42^. Each subunit contains a “bait region”, consisting of approximately 25 amino acids with many protease cleavage sites, responsible for the incredibly diverse range of α_2_M-interacting proteases^43^. Upon proteolysis of the bait region by active proteases, α_2_M becomes activated (α_2_M*) and undergoes a conformational change that results in covalent trapping of proteases within a steric cage^42^. This exposes the LRP1-binding domains^43^. It is well-established that α_2_M* is exclusively recognized by LRP1 within the LDL receptor family and undergoes clathrin-mediated endocytosis^42,44^.

It has been shown that stimulation of LRP1 with ligands such as α_2_M* or insulin induces LRP1 sorting to the PM by regulated exocytosis of intracellular vesicles, potentially LRP1 storage vesicles, rather than the classical and well-described Rab11-dependent recycling pathway, and that the receptor does not localize to acidic compartments^44^. As for ligands, α_2_M* follows a lysosomal degradation pathway, whereas apolipoprotein E and the heat shock protein gp96 avoid intracellular degradation and are instead exocytosed ^44^. Previously, our group demonstrated a high level of basal autophagy in human erythroleukemic cells (K562) and that inducing autophagy or overexpressing LC3 (an autophagy marker) promotes amphisome formation by mediating the fusion between MVBs with autophagosomes, inhibiting exosomal release^45^. In addition, we have shown that hemin (a physiological stimulant of erythroid maturation) can induce mitophagy (a type of autophagy) in K562 cells through the LRP1 receptor^32^.

Since the induction of autophagy is an indispensable event during the process of erythroid differentiation and that interaction between hemin and LRP1 induces autophagy in erythroleukemic cells, we aimed to study whether α_2_M* which is a specific ligand of LRP1 and not an inducer of erythropoiesis, can stimulate the autophagic pathway in this cell type. Our results show that α_2_M* stimulates autophagy and mitophagy in K562 cells through its binding to the LRP1 receptor, and that in the absence of both the receptor and the ligand, autophagic levels remain basal. In addition, we have verified the presence of LC3 in extracellular vesicles, so we believe that the interaction of the receptor with α_2_M* stimulates the fusion of MVBs with autophagosomes, forming amphisomes, and that these amphisomes subsequently fuse with the PM, releasing these LC3-positive vesicles.

## Materials and Methods

### Cell culture

K562 cells were cultured in RPMI 1640 Gibco (Invitrogen Argentina S.A) supplemented with 10% fetal bovine serum (The Cell Culture Company GBO), streptomycin (50 μg/ml) and penicillin (50 U/ml). UT7-EPO cells were maintained in MEM + GlutaMAXTM medium (ThermoFischer, France), supplemented with 10 % fetal bovine serum, 2mM L-glutamine, 1 % antibiotics penicillin/streptomycin and 2U/mL EPO (Sigma–Aldrich). Both cell lines were incubated at 37°C in a humidified atmosphere with 5 % CO2 and 95 % air.

### Reagents

Mowiol (Calbiochem 475,904-100 GM), Nitrocellulose membrane (GVS 1,213,314), Paraformaldehyde (Cicarelli 1,088,211), Penicillin/Streptomycin (Thermo Fisher Scientific 15,140,122), Potassium Chloride (Biopack, 7447-40-7), Potassium Phosphate Diacid (Tetrahedron), Saponin (Dalton), Sodium Bicarbonate (Anedra, 144-55-8), Sodium Chloride (Anedra, 7647-14-5), Sodium Phosphate Dibasic (Biopack, 7558-79-4), Tris-Chloride buffer (Promega, H5121), Chloroquine diphosphate salt (Cq) (Sigma–Aldrich) and Bafilomycin A1 (Baf) (Sigma–Aldrich).

### α-_2_-Macroglobulin Purification

α_2_M was purified from human plasma following a procedure at Centro de Investigación en Medicina Traslacional Dr. Severo Amuchástegui (CIMETSA) of the Instituto Universitario de Investigaciones en Ciencias Biomédicas Córdoba (IUCBC), U.A. CONICET, Córdoba, Argentina, following a procedure previously described^46^. The native form of α_2_M demonstrates proteinase inhibitory activity, but it is not recognized by LRP-1^47^. The activated form (α_2_M*) was generated by incubating α_2_M with 200 mM methylamine-HCl for 6 hours at pH 8.2, as previously described^48^.

### Antibodies

For Western blot (WB) and immunofluorescence, we employed: rabbit anti-LRP1β (Sigma–Aldrich, Canada), mouse anti-LRP1 (Abcam, Cambridge, MA), rabbit anti-LC3B (Sigma–Aldrich), mouse anti-β-actin (Sigma–Aldrich), rabbit anti-tubulin (Santa Cruz, CA), mouse anti-flotillin-2 (Santa Cruz, CA), rabbit anti-calnexin (Sigma Aldrich), anti-rabbit HRP (Jackson Lab), mouse anti-HRP (Jackson Lab), Alexa Fluor 488 anti-mouse (Sigma–Aldrich), Cy3 anti-rabbit (Sigma–Aldrich), Cy3 anti-mouse (Sigma–Aldrich), and Alexa Fluor 594 anti-rabbit (Sigma–Aldrich).

### Plasmids

YFP-Mito was a generous gift from Dr. Ernesto J. Podesta (School of Medicine, University of Buenos Aires-CONICET). GFP-Vector and RFP-LC3 were generated as indicated in our previous publications ^49,50^, GFP-Rab7 and GFP-Rab11 were kindly donated by Dr. María Isabel Colombo (Instituto de Histología y Embriología, CONICET, Facultad de Ciencias Médicas, Universidad Nacional de Cuyo, Mendoza, Argentina). GFP-CD63 was kindly donated by Dr. Ignacio Cebrian (Instituto de Histología y Embriología, CONICET, Facultad de Ciencias Médicas, Universidad Nacional de Cuyo, Mendoza, Argentina), GFP-CRISPR-LRP1 (pSpCas9(BB)-2A-GFP) and CRISPR-LRP1-puromycin (pSpCas9(BB)-2A-Puro) were produced in Dr. Thierry Galli’s laboratory in Paris, France.

### Plasmid transfection

A total of 2×10^6^ cells were used for each transient transfection per nucleoporation. Cells were centrifuged at low speed 800-1000 rpm for 4-5 min and the pellet was carefully resuspended in 100 μl of sterile 1X PBS (tempered to 37°C), with 1-2 μg of DNA which was previously added. The cells were homogenized and placed in 0.2 cm nucleofection cuvettes and transferred to the Amaxa Nucleofector II (IHEM, Mendoza, Argentina), as indicated by the manufacturer. Cells are then resuspended in RPMI 10% SFB medium and incubated at 37°C for recovery for at least 24 hours. After the recovery time, the percentage of transfected cells is quantified by fluorescence microscopy and can be used for subsequent assays. The nucleofector software has factory-loaded electrical standards for each cell line. In our case we used the K562 ATCC high efficiency protocol.

### Transduction

K562 and UT7 cells were transduced with lentiviral vector MISSION® shRNA (Sigma, CSTVRS Custom Lentiviral Particles) pLKO.1-CMV.tGFP, containing a short hairpin RNA scramble (shSCR) or a short hairpin RNA targeting LRP1 (shLRP1). In exponential phase cells were transducted by spinoculation, centrifugated at 800g for 30 min at 32°C with an MOI of 20. Transduction efficiency was evaluated by the percentage of GFP+ cells after 72 hours. GFP+ cells were sorted using the SONY SH800 cell sorter, and stable shSCR and shLRP1 cell lines were selected based on their transcriptional and protein expression of LRP1.

### SDS/PAGE and WB

After stimulation, cells were washed with PBS 1X and then lysed with a solution containing Triton X-100 and protease and phosphatase inhibitors. Total protein was quantified using the Pierce™ BCA Protein Assay Kit (Thermo Fisher scientific). For WB, we used 100 µg of total protein on 10–15% of SDS/PAGE, which was then transferred to a nitrocellulose membrane (GVS NitroPure). Blocking was performed for 1 hour with 5% fat-free milk in a 0.1% Tween 20 PBS solution. After two washes with PBS 1X, the primary antibodies were incubated overnight and peroxidase–conjugated secondary antibodies were incubated for 2 hours at room temperature. The corresponding bands were detected using the Pierce™ ECL Western Blotting Substrate kit (Thermo Scientific, 32209). Images were acquired using the ImageQuant LAS 4000 chemiluminescent imaging system and analyzed using ImageJ software.

### Trypan Blue cell viability assay

K562 cells were transfected with RFP-LC3 and GFPvector or GFP-CRISPR-LRP1, then incubated in the presence/absence α_2_M* [60 nM] for 24 hours and 2 hours in the presence/absence of Baf [100 nM] or Cq [50 µM]. After incubation, cells were centrifuged and washed with PBS 1X at 37°C to remove all the medium. Trypan Blue was added following the manufacturer’s guidelines. The percentage of cell death was obtained by analyzing the number of Trypan Blue dyed cells by the total cell number, counted (n=1000) in triplicate for each independent assay (n=3).

### Indirect immunofluorescence

For cell fixation, we used 4% paraformaldehyde for 15 min. Autofluorescence was blocked for 30 min with 50 mM NH4Cl in PBS, and cells were permeabilized for 20 min with 0.05% saponin in PBS containing 0.2% BSA. Primary antibodies were incubated for 2 hours and secondary antibodies for 1 hour at room temperature. Antibodies were prepared in PBS solution containing 1% BSA. Cells were mounted on glass slides using Mowiol with 0.1% Hoechst and examined with an Olympus FV-1000 fluorescence microscope. Images were analyzed by ImageJ software.

### Fluorescence microscopy

GFPv, GFP-CRISPR-LRP1, RFP-LCR or YFP-mito transfected K562 cells were incubated 24 hours with α_2_M* and 2 hours with Baf and subsequently analyzed *in-vivo* using an Olympus FV- 1000 Confocal Optical Microscope (IHEM, Mendoza, Argentina). Additionally, a 60X objective lens and 2X digital zoom (when necessary) were used for image acquisition. Then, 1024 × 1024 or 500 × 500 pixel RGB images were processed using ImageJ or FIJI free software for particle analysis and for colocalization analyses.

### Electron microscopy

K562 cells were incubated for 24 hours with α_2_M* and with Baf for 2 hours. Cells were then processed for electron microscopy as previously described ^51^. Briefly, the samples were fixed in 2,5% glutaraldehyde in 0.15 M sodium cacodylate buffer for 60 min, pre-infiltrated by adding a 2% aqueous solution of low-melting-point agarose and fixed using a 2% aqueous solution of osmium tetroxide (OsO₄) for 2 hours at room temperature. The samples were dehydrated using aqueous solutions with increasing concentrations of acetone, and the cells were then embedded in low-viscosity Spurr resin. Finally, 60–70 nm ultrathin sections were prepared using a Leica Ultracut R ultramicrotome and stained with aqueous solutions of lead citrate (0.5%) and uranyl acetate (2%) ^52^. The samples were observed under a Zeiss 900 electron microscope.

### Isolation of EVs and Nanoparticle Tracking Analysis

EVs were collected from 15 mL of culture medium containing K562 cells cultured for 24 hours with or without α_2_M*. To do this, the culture medium was placed on ice and centrifuged at 800 g for 10 minutes to sediment the cells and then centrifuged at 12,000 g for 20 minutes to remove cellular debris. The EVs were separated from the supernatant by centrifugation at 100,000 g for 2 hours. The EVs pellet was washed once in a large volume of PBS and finally resuspended in 100 ml of PBS (EV fraction). The samples were then analyzed using a nanoparticle tracking analyzer (NTA) (ZetaView PMX-120 - Particle Metrix), which allows for counting, measuring the size, zeta potential, and, in some cases, the fluorescence of the particles present in the analyzed material. The instrument was set to 22 °C, with a sensitivity of 75 and a shutter speed of 75. In scatter mode, measurements were taken at 11 different positions (three cycles per position) at a rate of 30 frames per second.

### Image processing

Briefly, the puncta quantification was made using the following protocol: First, RGB images were converted to 8-bit images, and color-channels were separated. Then, Brightness and Contrast were manually adjusted using the HiLo LUT conversion table. After this, the image was duplicated and a manual threshold was applied, followed by the selection of the specific region of interest (ROI). Finally, ImageJ’s particle analysis was used to automatically obtain the number of positive structures for different targets inside the ROI. For colocalization analysis, the ImageJ JACoP plugin was used. The same image processing mentioned above was used, with the addition of a mask creation using the specific ROI before applying the plugin. The program calculates the overlap of pixels from each fluorophore emission channel and generates an image showing the channel overlap in RGB color, along with the percentage of colocalization based on a randomness metric, calculated using Pearson’s and Manders’ statistical methods.

### Statistical analysis

Statistical analyses were performed using Prism 8 (GraphPad Software). Data were evaluated using Mann-Whitney or Kruskal-Wallis tests, and results are presented as the mean ± SEM from at least two independent experiments. Comparisons were performed using ANOVA along with Tukey and Dunnett tests. Significant differences: *P < 0.01; **P < 0.005.

## Results

### Activated α_2_M stimulates autophagy in erythroleukemic cells

We previously demonstrated that LRP1 is necessary to promote hemin-stimulated autophagy in human erythroid-like cells especially the K562 and HEL92.1.7 cell line^53^. This prompted us to study if α_2_M* is also able to activate autophagy in K562 cells. For this purpose, K562 cells were cultured in the presence or absence of α_2_M* and with Baf or Cq to hamper autophagosome-lysosome interaction (Figure 1). Subsequently, the levels of the LRP1 and LC3 proteins were analyzed by Western blot, and this analysis was supplemented with indirect immunofluorescence studies and confocal microscopy. As we can observe in Figure 1, A-B, there were no significant differences in the amount of LRP1 protein between different conditions. On the other hand, examination of LC3 showed that the lipidated form LC3-II accumulated when autophagic flow was blocked with Cq compared to the control. Interestingly, this rise was significantly greater in cells treated with α_2_M* combined with Cq than in cells treated with Cq only, suggesting an increase in autophagy (Figure 1, A and C). These results were confirmed by indirect immunofluorescence, in which endogenous LC3 and LRP1 were analyzed under each experimental condition (Figure 1, D). Quantitative analysis of LRP1-positive fluorescent dots showed no statistically significant variations (Figure 1, E), whereas the number of LC3-positive vesicles per cell increased significantly in cells treated with α_2_M* and Cq (Figure 1, F). Taken together, these results demonstrate that treatment with α_2_M* increases autophagic activity in K562 cells.

**Figure 1:**
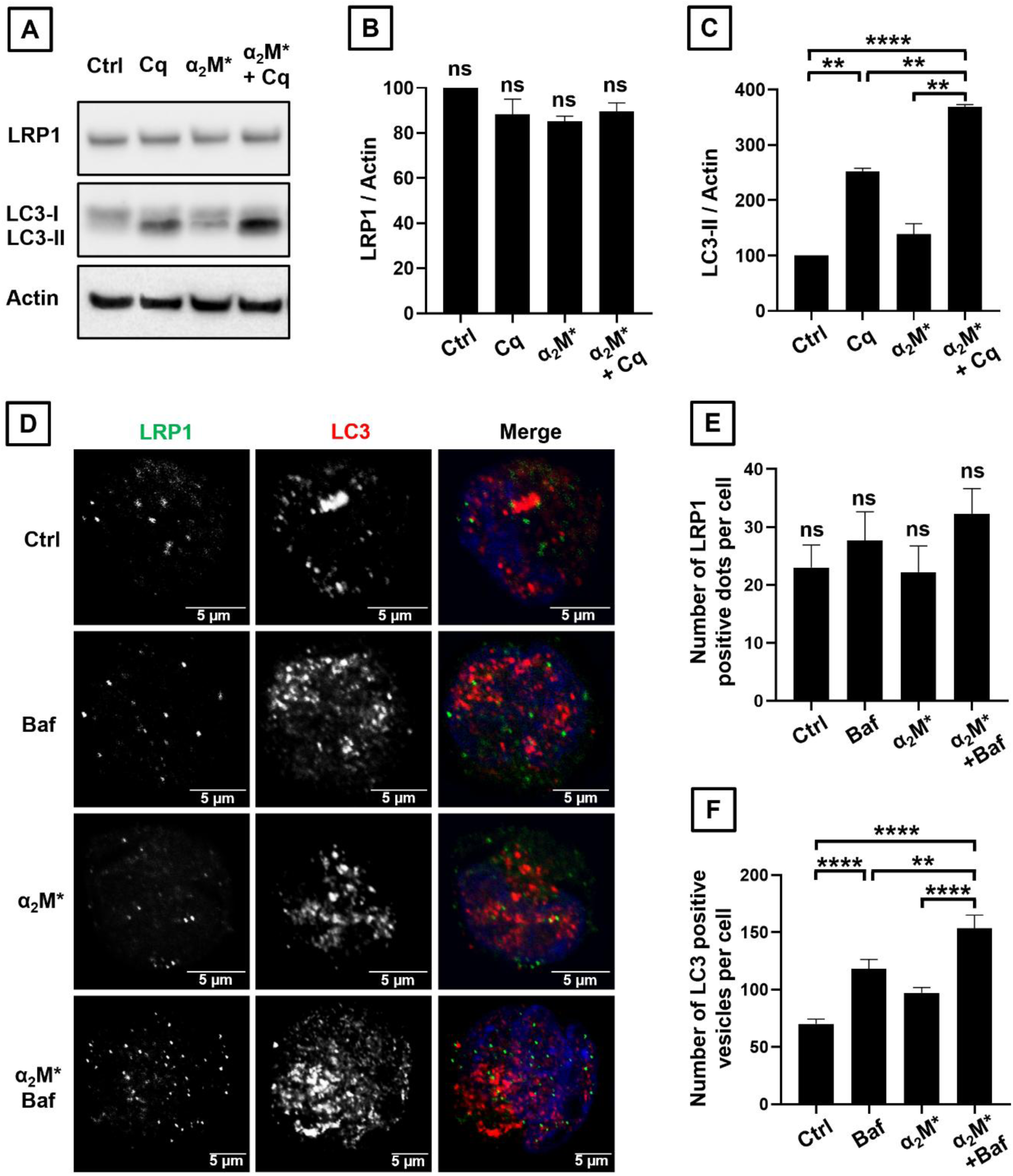
α_2_M* increases LC3-II protein levels in K562 cells. **(A)** Western blot showing LRP1 and LC3-II protein levels in K562 cells cultured 24 hours in absence (Ctrl) or presence of activated α-_2_-Macroglobulin (α_2_M*) [60 nM]. Cq [50 µM] was added for 2 hours to block autophagic flow. Rabbit anti-LRP1, rabbit anti-LC3, and, as a loading control, mouse anti-β-actin antibodies were used. **(B)** Quantification of LRP1/actin and **(C)** LC3-II/actin protein levels. **(D)** Confocal microscopy images of K562 cells cultured under the same conditions, replacing Cq with Baf [100 nM] and immunolabeled with the primary antibodies mouse anti-LRP1 and rabbit anti-LC3. Secondary antibodies anti-mouse Alexa 488 and anti-rabbit Cy3 were used, respectively. Nuclei were stained with Hoechst. Scale bar: 5 µm. **(E)** Quantification using ImageJ-Fiji of the number of LRP1-positive dots or **(F)** LC3-positive dots per cell. The bars represent the mean of three independent experiments ± SE. ** = p < 0.01 and **** = p < 0.0001.

### α_2_M* induces autophagy in K562 cell line through the LRP1 receptor

Since we previously demonstrated that α_2_M* stimulates autophagy in erythroleukemic cells, we were interested in determining the role of LRP1 in this context. To this end, K562 and UT7 cell lines were transduced with lentiviral vectors containing either a short-hairpin scramble RNA (shSCR) or one targeting LRP1 (shLRP1), and wild-type (WT) cells were used as a control. The cells were then incubated for 24 hours in the presence or absence (Ctrl) of α_2_M*, and Cq was added for 2 hours to hamper autophagosome-lysosome interaction. Subsequently, LRP1 and LC3 levels were analyzed by Western blot (Figure 2). The analysis showed a marked decrease in LRP1 levels in K562 (Figure 2, A–B) and UT7 (Figure 2, D–E) cells transfected with shRNA specific to the receptor, compared to WT and shSCR cells, confirming the efficiency of silencing. Afterwards, we assessed LC3-II levels and observed that α_2_M* increased autophagy in K562 WT and UT7 shSCR cells as expected, whereas in shLRP1 cells this increase was significantly reduced to the baseline level of autophagy for these cells (Figure 2, C and F). These results indicate that the induction of autophagy by α_2_M* requires the expression of the LRP1 receptor in both K562 and UT7 cells.

**Figure 2:**
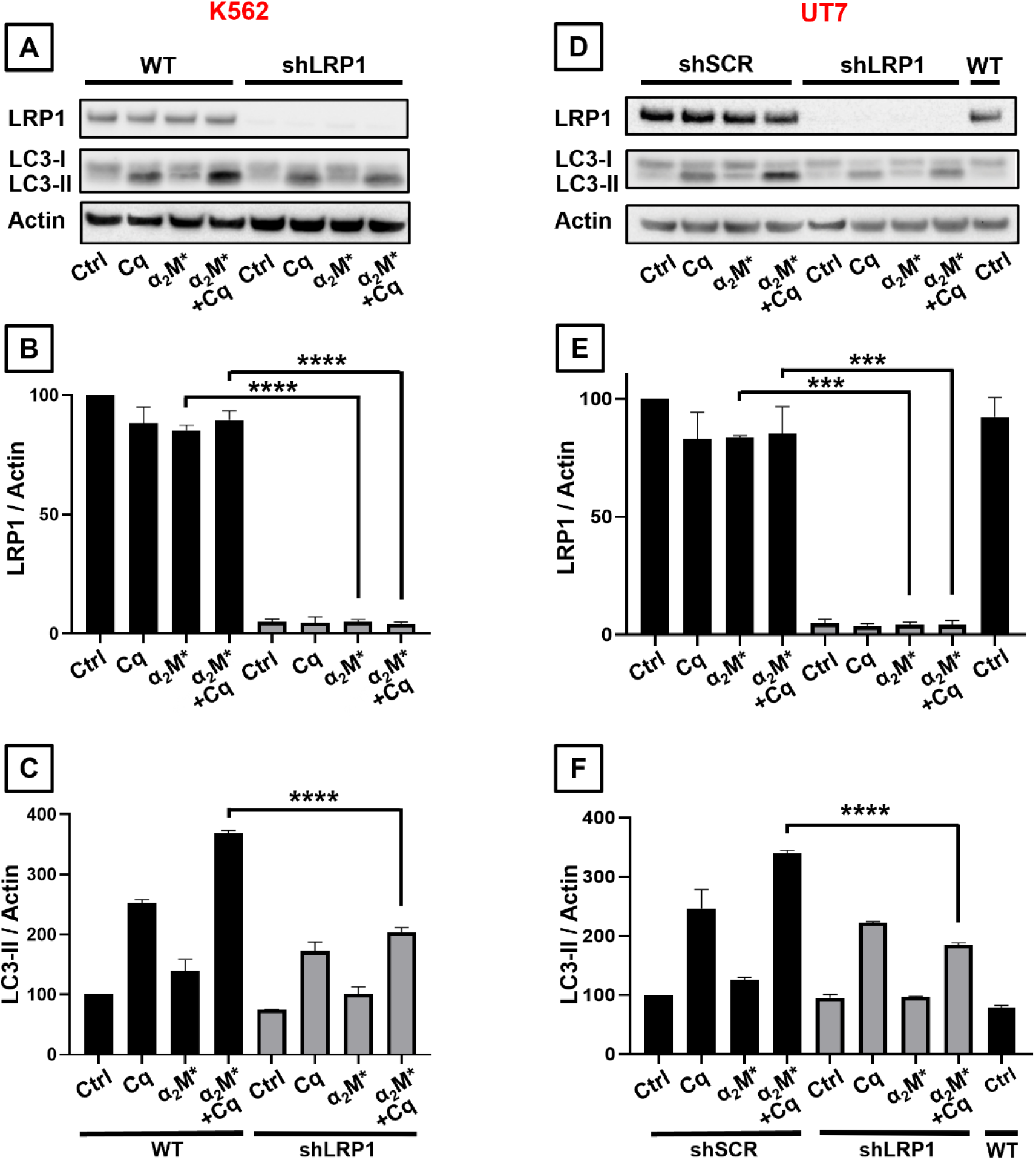
The α_2_M*-LRP1 interaction induces the autophagy pathway in K562 and UT7 cell lines. **(A)** WB analysis of WT and shLRP1 K562 cells incubated in the presence or absence (Ctrl) of Cq [50 µM] and/or α_2_M* [60 nM]. Rabbit anti-LRP1, rabbit anti-LC3, and mouse anti-actin primary antibodies were used, along with rabbit anti-HRP and mouse anti-HRP secondary antibodies. **(B)** Quantification of LRP1 and LC3 protein levels **(C)** in WT and shLRP1 K562 cells. **(D)** Western blot for UT7 WT, shSCR, and shLRP1 cells incubated under the same conditions as the K562 cells. **(E)** Quantification of LRP1 and LC3 protein levels **(F)** in WT, shSCR, and shLRP1 UT7 cells. Bars represent the mean of three independent experiments ± SE. *** = p < 0.001 and **** = p < 0.0001. **(C)** Percentage of cell viability after transfection, determined by the trypan blue exclusion assay. 10% DMSO and high temperatures were used as negative controls. **(D)** Quantification of the number of RFP-LC3-positive vesicles per cell, under each condition, using ImageJ-Fiji. Bars represent the mean of three independent experiments ± SE. ** = p < 0.01 and **** = p < 0.0001.

Furthermore, these findings were validated by generating LRP1-knockout K562 cells using the CRISPR/Cas9 machinery (Figure 3). For this purpose, the cells were co-transfected with GFP-CRISPR-LRP1 construct and RFP-LC3, using cells co-transfected with the GFP-vector and RFP-LC3 as a control. Following transfection, the cells were incubated for 24 hours in the presence or absence of α_2_M*, and Baf was added during the final 2 hours to hamper autophagosome-lysosome interaction (Figure 3, A). The efficiency of the LRP1 knockout was assessed by Western blot, analyzing LRP1 levels in WT K562 cells and in cells transfected with GFP-CRISPR-LRP1 or with a second CRISPR construct targeting LRP1 that contains a puromycin resistance gene, using actin as a loading control (Figure 3, B). Cell viability analysis using the Trypan blue exclusion assay showed that LRP1 knockout did not affect cell viability under the experimental conditions analyzed (Figure 3, C). Subsequently, the number of LC3-positive vesicles per cell was quantified by confocal microscopy. In cells transfected with GFP-vector, incubation with α_2_M* induced an increase in the number of LC3-positive vesicles (Figure 3, D), comparable to that previously observed in WT cells (Figure 1, A and C). In contrast, in cells transfected with GFP-CRISPR-LRP1, treatment with α_2_M* did not result in a significant increase in the number of LC3-positive vesicles; instead, a marked reduction was observed compared to control conditions (cells transfected with GFP-vector) (Figure 3, D). Taken together, these results confirm that α_2_M*-induced autophagic activation depends on the presence of the LRP1 receptor in erythroleukemic cells.

**Figure 3:**
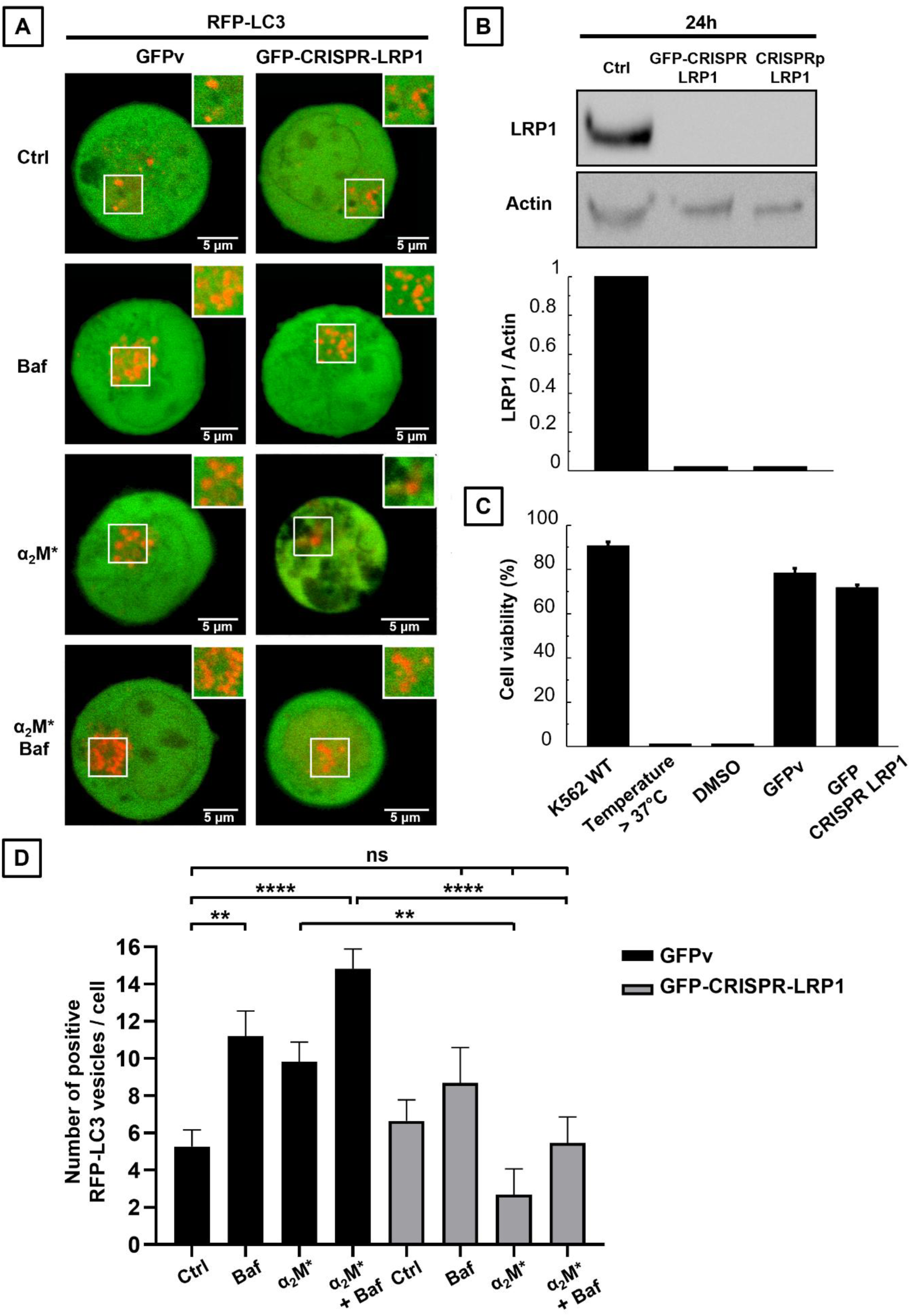
α_2_M*-induced autophagy activation depends on the LRP1 receptor in K562 cells. **(A)** *In vivo* confocal microscopy images of K562 cells cotransfected with RFP-LC3 and GFPvector (GFPv) or GFP-CRISPR-LRP1. 24 hours post-transfection, the cells were incubated in the absence (Ctrl) or presence of α_2_M* [60 nM] for 24 hours. In the last 2 hours, Baf [100 nM] was added to block autophagic flow. **(B)** WB showing LRP1 levels in WT K562 cells or cells transfected with GFP-CRISPR-LRP1 or CRISPRp-LRP1. Actin was used as a loading control. **(C)** Percentage of cell viability after transfection, determined by the trypan blue exclusion assay. 10% DMSO and high temperatures were used as negative controls. **(D)** Quantification of the number of RFP-LC3-positive vesicles per cell, under each condition, using ImageJ-Fiji. Bars represent the mean of three independent experiments ± SE. ** = p < 0.01 and **** = p < 0.0001.

### Activated alpha-_2_-Macroglobulin stimulates mitophagy in erythroleukemic cells

Given the central role of mitophagy during erythroid maturation, we investigated whether α_2_M* induces this process in erythroleukemic cells (Figure 4). For this purpose, K562 cells were cotransfected with YFP-Mito, as a mitochondrial marker, and RFP-LC3, as a marker for autophagosomes. The cells were then incubated for 24 hours in the absence (Ctrl) or presence of α_2_M*; Baf was added during the last 2 hours to allow for the accumulation of autophagic structures, and the cells were analyzed *in vivo* by confocal microscopy (Figure 4, A). The colocalization of the red signal and the yellow-green signal and vice versa was then assessed (Figure 4, B and C). Results showed that treatment with α_2_M* induced an increase in colocalization between LC3 and mitochondria, evidenced by the overlap of the fluorescent signals and enhanced by blocking autophagic flux, suggesting activation of the mitophagy process in these cells. Moreover, to confirm that the selective degradation of mitochondria by autophagy increases in the presence of α_2_M*, we assessed mitophagy at the ultrastructural level using transmission electron microscopy under the same incubation conditions. Electron microscopy analysis revealed the presence of autophagosomes under all conditions; and in some cases, mitochondria were identified within these structures (Figure 4, D: red arrows). Furthermore, consistent with the results obtained previously, quantification of the number of autophagosomes per cell showed a significant increase in those treated with α_2_M*, an effect that was enhanced by blocking autophagic flux (Figure 4, E). Therefore, these results confirm that treatment with α_2_M* increases mitophagy in K562 cells, as evidenced at the ultrastructural level.

**Figure 4:**
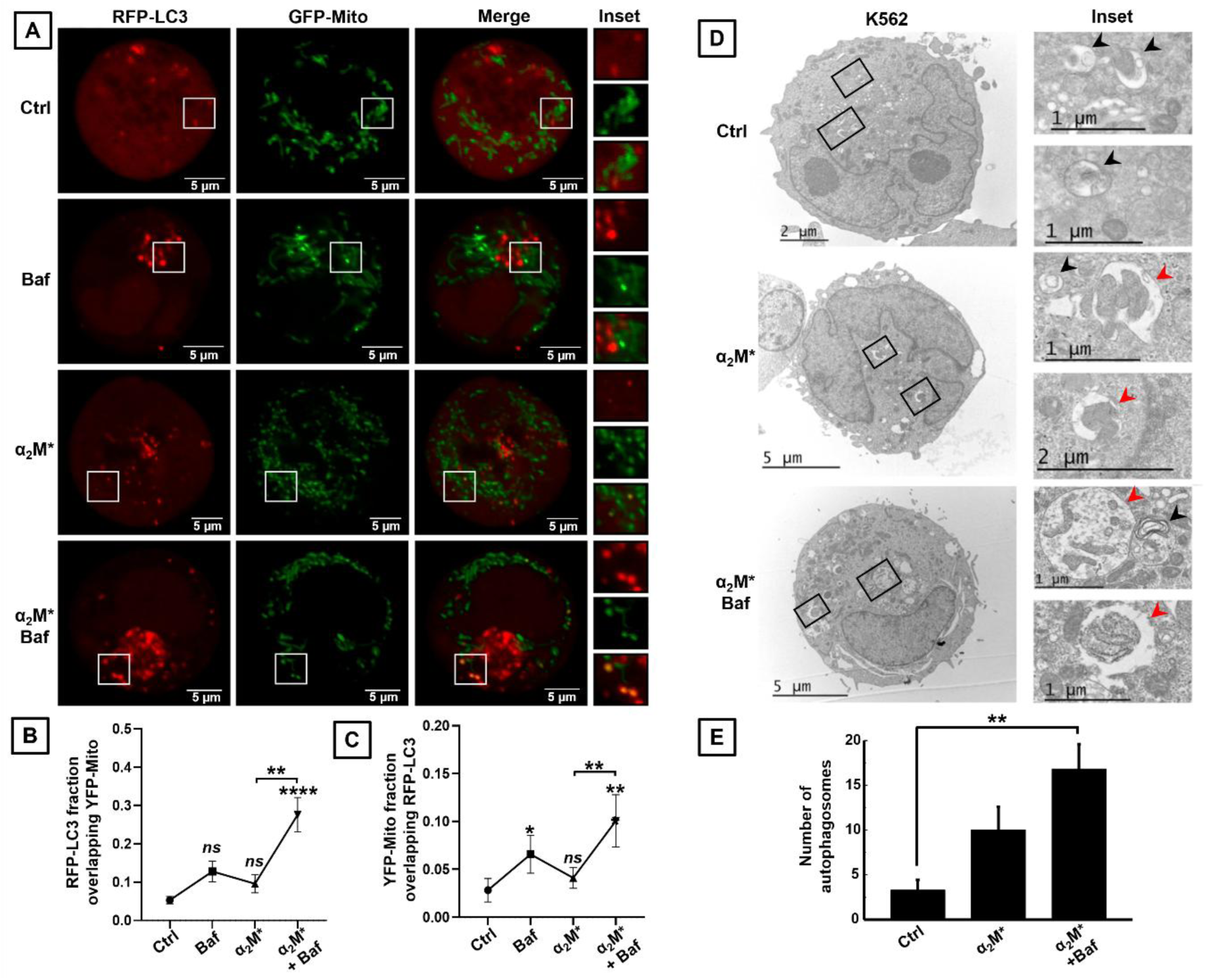
α_2_M* induces mitophagy in the K562 cell line. **(A)** *In vivo* confocal microscopy images of K562 cells cotransfected with YFP-Mito and RFP-LC3 and incubated in the absence (Ctrl) or presence of α_2_M* [60 nM] for 24 hours and Baf [100 nM] during the last 2 hours to block autophagy. **(B)** Manders colocalization of the RFP-LC3 fraction with YFP-Mito and of the YFP-Mito fraction with RFP-LC3 **(C). (D)** Electron microscopy images of K562 cells incubated in the absence (Ctrl) or presence of α_2_M* [60 nM] for 24 hours and 2 hours with Baf [100 nM] to block autophagic flow. Black arrows indicate autophagosomes and red arrows indicate autophagosomes containing mitochondria. **(E)** Quantification of the number of autophagosomes in each condition. Results represent the mean of three independent experiments ± SE. ** = p < 0.01, *** = p < 0.001, and **** = p < 0.0001.

### Interactions of Autophagosomes with Endocytic and Recycling Compartments

Due to the intrinsic relationship between autophagosome formation and the endocytic track during erythroid clearance, we investigated the potential intersection of these pathways in our experimental model. Specifically, we focused on the link between autophagosomes, multivesicular bodies, and recycling endosomes, as the formation of hybrid vesicles like amphisomes suggests a highly integrated network that coordinates both the degradation of internal components and communication with the extracellular environment through exosomes (Figure 5). To this end, we analyzed K562 cells cotransfected with RFP-LC3 and GFP-CD63, a classic marker of late endosomes and MVBs, using *in vivo* confocal microscopy under basal conditions and in the presence of α_2_M* (Figure 5, A). We then measured the colocalization of the green signal (late endosomes/MVBs) with the red signal (autophagosomes) and vice versa (Figure 5, B and C). The analysis revealed a significant increase in the association between LC3 and CD63 in the presence of α_2_M* compared to the control condition. These results indicate that α_2_M* increases the association between autophagosomes and late endosomes/MVBs in K562 cells. Subsequently, based on an analysis of the interaction between the autophagy pathway and late endosomal compartments, we investigated the possible relationship between LC3 and recycling endosomes. To this end, K562 cells were cotransfected with RFP-LC3 and GFP-RAB11 and analyzed by *in vivo* confocal microscopy in the absence or presence of α_2_M* (Figure 5, D). In contrast to the significant increase observed for CD63, the colocalization between LC3 and RAB11 was not significant (Figure 5, E and F). It is worth noting that only a fraction of the LC3-positive vesicles colocalized with RAB11, which is to be expected given that RAB11 marks recycling compartments, many of which do not participate in the formation of autophagy-associated amphisomes or MVBs. In this context, the observed increase in α_2_M* levels suggests that a subset of RAB11-containing vesicles may be recruited into the autophagic pathway.

**Figure 5:**
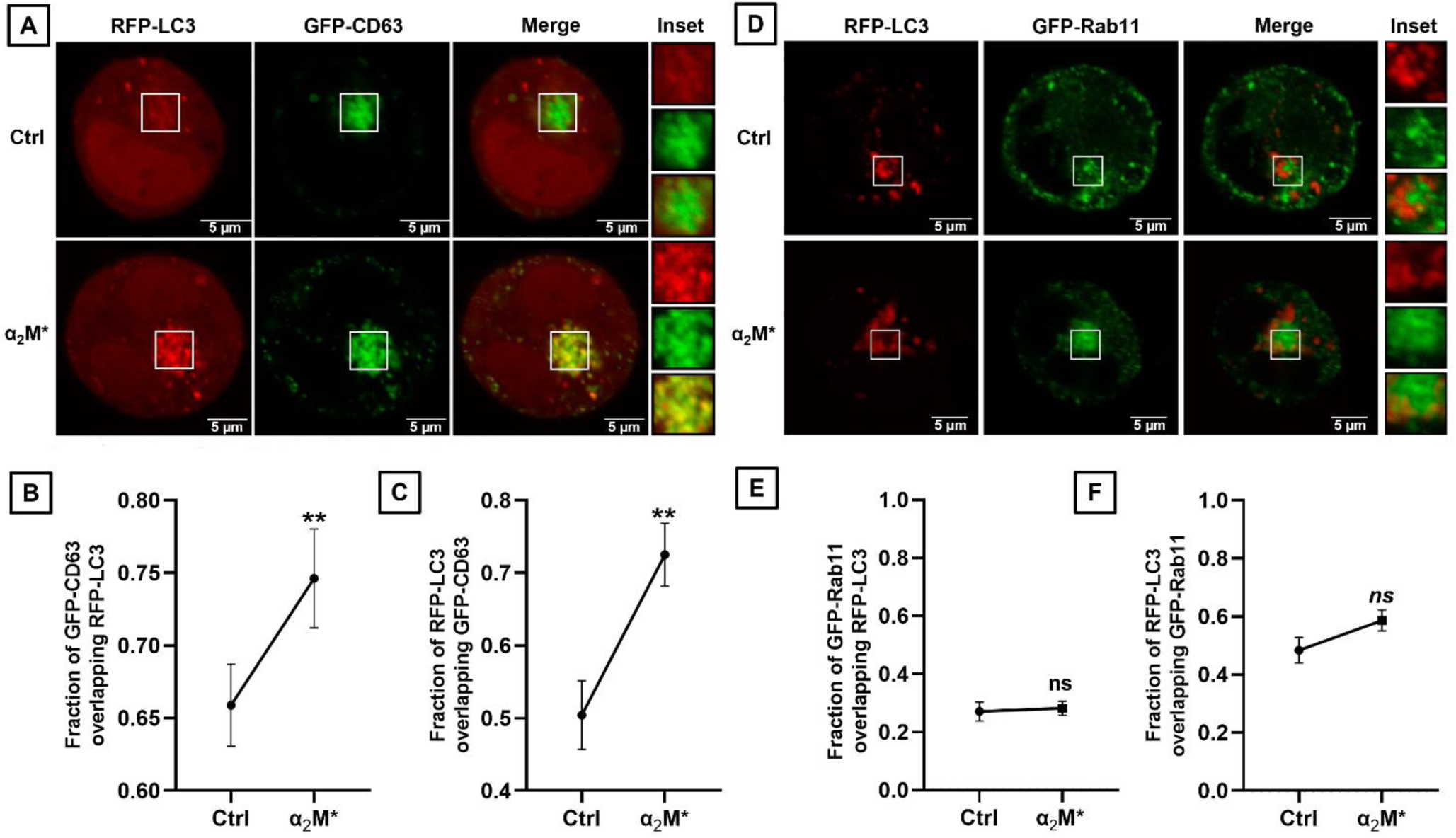
The association between autophagosomes and late endosomes/MVBs increases in the presence of α_2_M*. **(A)** *In vivo* confocal microscopy images of K562 cells cotransfected with RFP-LC3 and GFP-CD63 and incubated in the absence (Ctrl) or presence of α_2_M* [60 nM] for 24 hours. **(B)** Colocalization according to the Manders coefficient of the GFP-CD63 fraction with RFP-LC3 and **(C)** of the RFP-LC3 fraction with GFP-CD63. **(D)** *In vivo* confocal microscopy images of K562 cells cotransfected with RFP-LC3 and GFP-RAB11 and incubated in the same conditions as A. **(E)** Manders quantitative analysis of colocalization for the RAB11 signal colocalizing with LC3 and **(F)** the LC3 signal colocalizing with RAB11. Statistical comparisons were performed using the Mann–Whitney test, and results represent the mean of three independent experiments ± SE. ** = p < 0.01.

### Release of LC3-positive EVs from K562 cells in the presence of α_2_M*

To investigate the possible role of the LRP1 receptor in exosome release during erythroid maturation, we analyzed the population of EVs released by K562 cells cultured under control and α_2_M* conditions (Figure 6). The EVs released into the medium were purified by differential ultracentrifugation and characterized by NTA and Western blot as described in the methodology. NTA analysis of the size distribution of the EVs revealed, in both experimental conditions, a predominant peak within the size range consistent with exosomes (Figure 6, A). Furthermore, quantification of the percentage of released EVs showed a trend toward an increase in the α_2_M*-treated condition compared to the control (Figure 6, B). Subsequently, using Western blot, we observed that, under both conditions, the samples were negative for calnexin (an endoplasmic reticulum marker), ruling out contamination by cellular components, and positive for flotillin-2, a protein widely used as an exosome marker. Additionally, LC3 was detected in EVs in both the control and α_2_M* conditions (Figure 6, C). These results indicate that K562 cells release LC3-positive EVs that could be exosomes and that there is a tendency for an increase in these EVs in the presence of α_2_M*.

**Figure 6:**
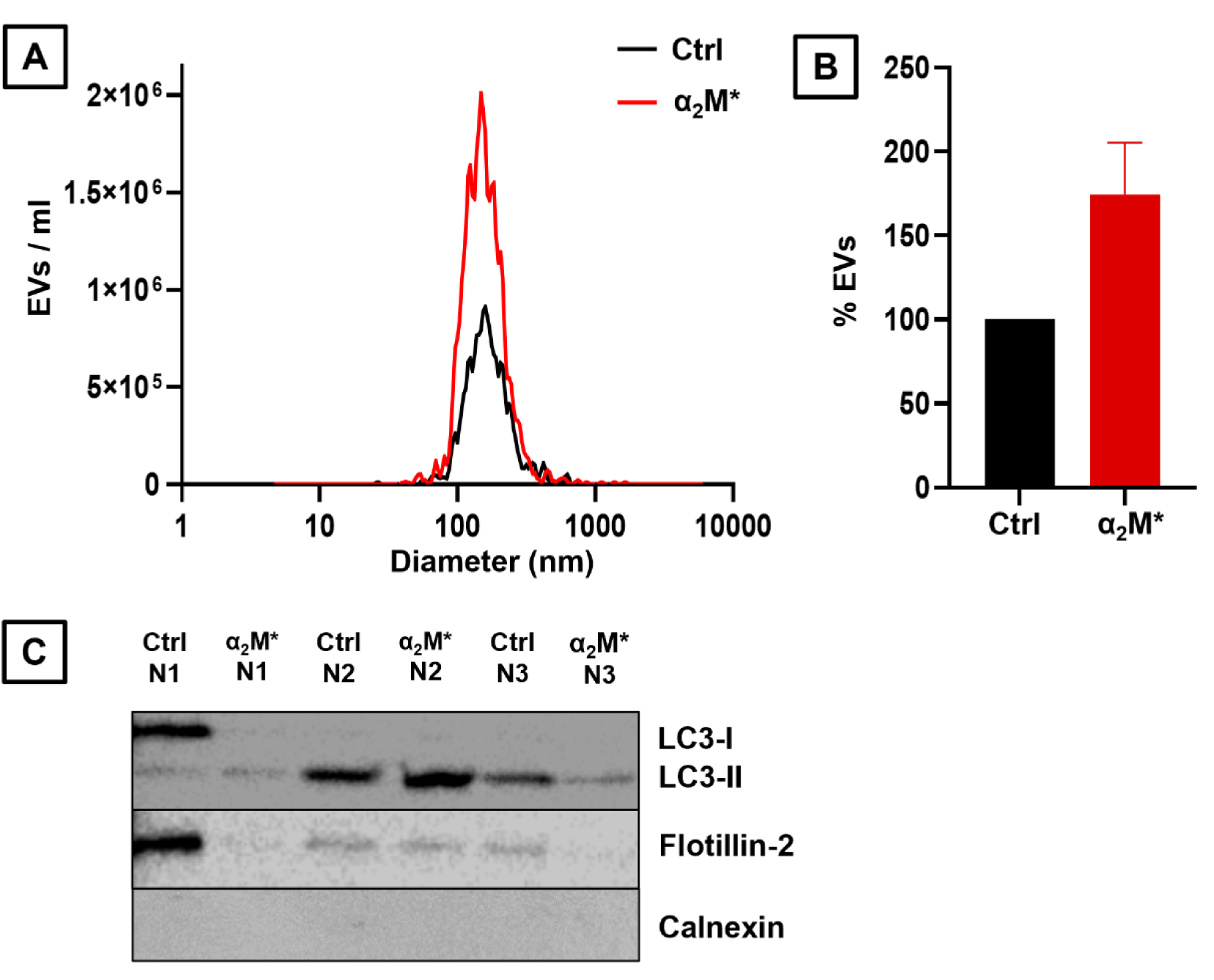
K562 cells release LC3-positive EVs. K562 cells were cultured in the absence (Ctrl) or presence of α_2_M* [60 nM] for 48 hours, and the EVs released into the medium were isolated by differential ultracentrifugation. **(A)** Analysis of EV size distribution by NTA (N=2). **(B)** Quantification of the percentage of EVs released under both conditions (N=2). **(C)** Western blot of EVs purified under both conditions (N=3). Rabbit anti-LC3, mouse anti-flotillin-2, and rabbit anti-calnexin primary antibodies were used, along with rabbit anti-HRP and mouse anti-HRP secondary antibodies.

## Discussion

The association of the receptor LRP1 with autophagy was first described by Dr. Yahiro, who established that LRP1 silencing blocks LC3-II accumulation and modulates autophagy and apoptosis induced by specific stimuli, such as the *Helicobacter pylori* VacA toxin^54^. Subsequently, our group provided additional evidence by showing that hemin, a physiological inducer of erythroid differentiation, activates autophagy in K562 cells, accompanied by an increase in LRP1 expression and its colocalization with autophagosomes^32^. Given that, our working hypothesis focused on determining whether activation of this receptor by its physiological ligands could trigger this process. Based on this premise, we evaluated whether α_2_M* acts as a specific modulator of cellular degradation pathways, which would allow us to identify an intrinsic control mechanism during erythroid maturation. We demonstrated that when K562 cells are incubated with α_2_M* in the presence of Baf or Cq, the levels of LC3-II increase significantly compared to cells treated with α_2_M* alone (Figure 1). This observation suggests that α_2_M* does not act as a simple ligand destined for lysosomal degradation, but rather as a stimulus capable of activating intracellular pathways that converge in the formation and maturation of autophagosomes.

To confirm that this autophagic induction was strictly dependent on the interaction between the ligand and its receptor, we next addressed whether the absence of LRP1 would abrogate the effects of α_2_M*. In this context, silencing the receptor results in a significant decrease in the autophagic response^32^. Consistent with these antecedents, our results show that silencing LRP1 in K562 and UT7 cells leads to a significant reduction in LC3 levels, particularly its lipidated form. Notably, this inhibition is observed under basal conditions and is even more pronounced in α_2_M\***-**stimulated cells, reinforcing the notion that LRP1 acts as a central regulator of ligand-dependent autophagic induction (Figure 2). The validation of these findings in the UT7 line is especially relevant, as it demonstrates that the inductive effect depends on the stimulus exerted by α_2_M* rather than intrinsic peculiarities of a specific cell line. The role of LRP1 as a mediator was further confirmed through the generation of LRP1-knockout K562 cells via CRISPR/Cas9, where the absence of the receptor completely abolished the α_2_M*-induced increase in LC3-II levels (Figure 3).

As a cellular remodeling mechanism, autophagy facilitates the selective removal of organelles during erythropoiesis, with mitophagy being a crucial event in the formation of functional erythrocytes^55^. This essential step in terminal differentiation reduces oxidative stress through the removal of mitochondria and ensures the metabolic adaptation necessary for the maturation of reticulocytes^56^. Given the central role of mitophagy during erythroid maturation, we investigated whether α_2_M* induces this process in erythroleukemic cells (Figure 4). In this regard, colocalization studies between mitochondria labeled with YFP-Mito and RFP-LC3 allowed for a global assessment of the association between both structures, revealing a significant increase in mitochondria recruited to autophagic structures following α_2_M* stimulation. These results were confirmed by Transmission Electron Microscopy (TEM), where stimulated K562 cells showed a high percentage of autophagosomes containing mitochondria. The mitochondrial ultrastructure is easily recognizable via TEM by the presence of the double membrane, the higher electrodensity of the matrix, and the characteristic arrangement of the cristae. Together, these observations confirm that α_2_M* stimulation not only induces autophagy but specifically promotes mitochondrial clearance in erythroleukemic cells.

Hematopoiesis, and particularly terminal erythropoiesis, involves intense cellular remodeling that requires the coordination of multiple intracellular traffic routes, including the endocytic and autophagic pathways^2,4^. The endocytic pathway is a central axis of traffic, allowing for the internalization, sorting, recycling, or degradation of membrane proteins and extracellular components through a dynamic network of endosomal compartments^57^. Furthermore, autophagy can connect with the endosomal pathway through hybrid structures called amphisomes^58^. Regarding LRP1, previous studies have shown that its binding to α_2_M* induces receptor internalization and redistribution toward specific endosomal compartments, from which LRP1 can be rapidly recycled to the plasma membrane via non-canonical, Rab11-independent pathways, while the ligand is directed toward acidic degradative complexes. This ligand-dependent traffic suggests that LRP1 activation regulates not only endocytosis but also the functional organization of the endosomal pathway^59^. In this work we observed, via *in vivo* confocal microscopy, a significant increase in colocalization between LC3 and CD63 in K562 cells in the presence of α_2_M* compared to control conditions (Figure 5). CD63 is a classic marker of MVBs and exosomes, used to identify late endosomal compartments involved in EVs biogenesis. In contrast, the low colocalization between LC3 and Rab11 suggests that the Rab11-dependent recycling route is not the primary pathway of interaction with autophagosomes under these conditions. This reinforces the idea that, during erythroid maturation, LC3-positive compartments are mainly associated with degradative or secretory routes linked to MVBs rather than recycling mechanisms.

In this context, the observed trend toward increased EVs release in the presence of α_2_M* suggests that LRP1 activation could modulate the fate of MVBs. Moreover, the detection of LC3 in EVs released by K562 cells provides evidence that components of the autophagic machinery can be secreted into the extracellular space (Figure 6). This finding is consistent with the concept of secretory autophagy, where LC3-II may participate in both amphisome formation and the selective packaging of cargo within intraluminal vesicles^60,61^. In a demanding system like erythropoiesis, this could represent a complementary pathway for cellular clearance, where the secretion of LC3-positive EVs alleviates the lysosomal load and contributes to plasma membrane remodeling. Integration of our experimental evidence demonstrates that α_2_M* stimulation triggers a functional reorganization of the endosomal pathway, favoring the interaction between autophagosomes and MVBs.

### Conclusions

Terminal erythropoiesis is characterized as a dynamic process of cellular reorganization, in which organelle remodeling, the reconfiguration of vesicular trafficking, and metabolic adaptations precisely align with the functional demands of each stage of differentiation. In this context, the results presented in this study demonstrate that stimulation with α_2_M* not only induces the formation of autophagosomes but also promotes their interaction with late endosomal compartments and MVBs, leading to hybrid structures (amphisomes) and the release of EVs containing components of the autophagic machinery such as the LC3 protein (Figure 7). From a functional perspective, this link between lysosomal degradation and vesicular secretion is particularly significant in the erythroid context, where the cell undergoes intense remodeling of membranes and organelles; it emerges as a flexible mechanism that would allow the fate of cellular material to be adjusted according to metabolic state and microenvironmental signals. Furthermore, the results obtained suggest that activation of LRP1 by a specific extracellular ligand, such as α_2_M*, is sufficient to induce a robust autophagic response in erythroid cells, independent of classic intracellular stimuli associated with differentiation, such as hemin. Furthermore, the release of LC3-positive EVs suggests the involvement of secretory autophagy mechanisms or alternative cellular clearance pathways, which opens the possibility that these vesicles contribute to intercellular communication in the bone marrow, modulating the erythroid niche or the response of neighboring cells. In this regard, alterations in the quantity or content of EVs could have functional consequences on hematopoietic homeostasis.

**Figure 7.**
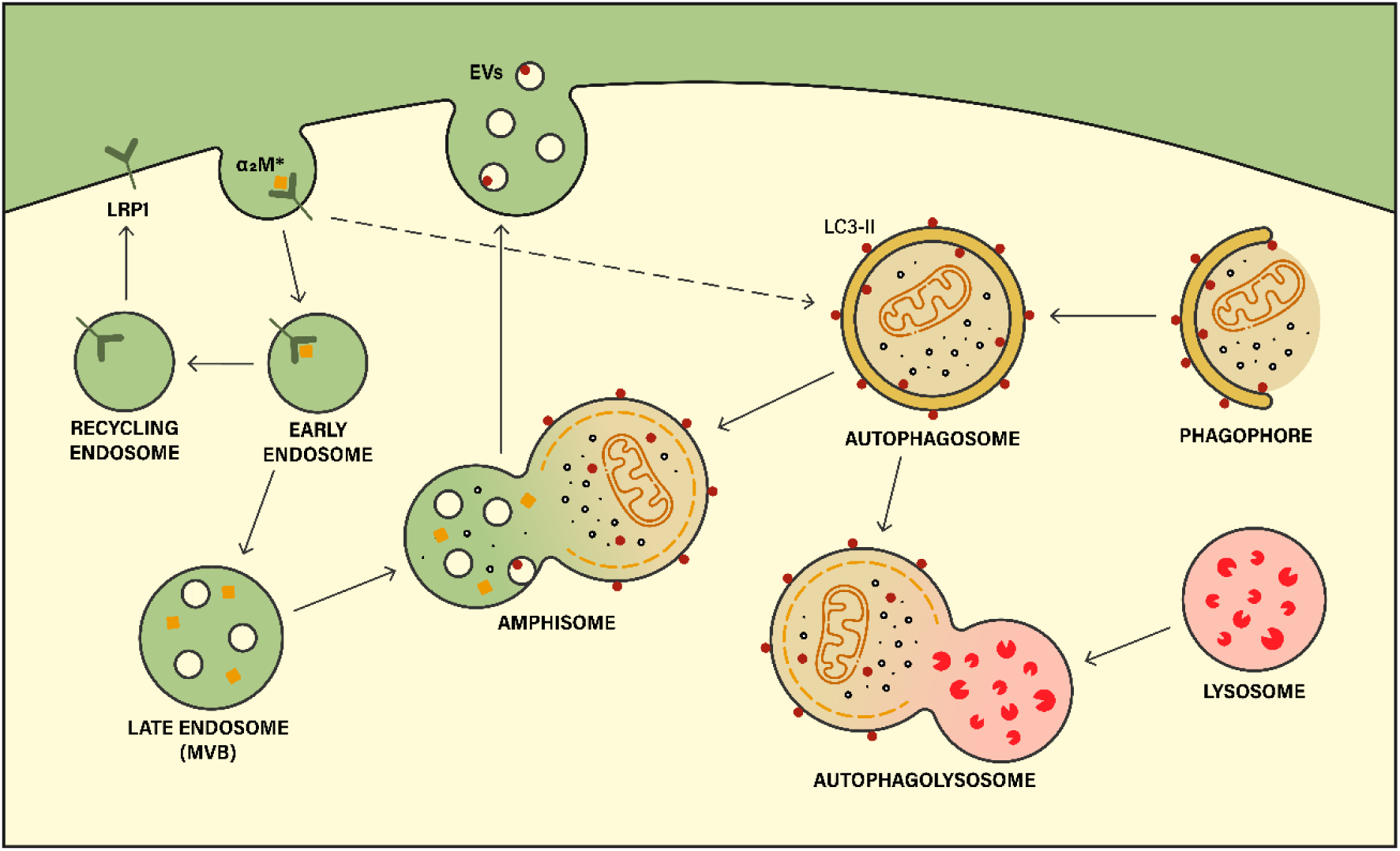
Proposed model. Based on the conclusions drawn from the results and in conjunction with the previous evidence, we propose a model in which the α_2_M*–LRP1 interaction in erythroid cells induces the formation of autophagosomes, which allows for the degradation of cellular components such as mitochondria via mitophagy. Meanwhile, at the same time, its interaction promotes the association between autophagosomes and late endosomal compartments or MVBs, giving rise to hybrid structures called amphisomes and the release of EVs containing components of the autophagic machinery, such as the LC3 protein, as an alternative pathway for cellular clearance and reorganization.

